# Liquid Crystal-like Self-Organization of Glioblastoma Prevents Cell Density Induced Migratory Arrest

**DOI:** 10.64898/2025.12.01.691550

**Authors:** Urszula Hohmann, Chalid Ghadban, Julian Prell, Christian Strauss, Faramarz Dehghani, Tim Hohmann

**Affiliations:** Department of Anatomy and Cell Biology, Medical Faculty, Martin Luther University Halle-Wittenberg, 06108 Halle (Saale), Germany; Department of Neurosurgery, Medical Faculty, Martin Luther University Halle-Wittenberg, 06120 Halle (Saale), Germany

**Keywords:** Cell migration, glioblastoma, liquid crystals, cell density, oncostreams, proliferation, collective migration

## Abstract

Glioblastoma is the most lethal and frequent type of primary brain tumors, characterized by a high proliferative and infiltrative capacity, leading to frequent recurrences after therapy. Here, we used live cell imaging to analyze the effect of cell density variations on the migratory capacity of established and primary glioblastoma cell lines. We found that proliferation events promoted local velocity of glioblastoma cells, but had only a small effect on large spatial scales. Furthermore, two phenotypes were found when subjecting glioblastoma cells to cell density gradient. One phenotype was characterized by the active migration of cells, independent of proliferation, while the other was mostly driven by cell proliferation. Lastly, the analysis of effects of an overall increasing cell density demonstrated that cells showing signs of self-organization, forming liquid crystal-like structures, are able to escape the expected cell-density induced migratory arrest. Notably, the emergence of small-scale liquid crystal-like order was associated with a promotion of cellular migration, even in cell populations that were largely in a state of migratory arrest. Thus, structure formation could help glioblastoma cells to move efficiently in states of high confinement, maintaining infiltrative properties.

## Introduction

Glioblastoma (GBM) belong to a class of highly aggressive tumors, with a median survival of approximately 14 month (1). Amongst others GBM are characterized by a strong proliferation and infiltration into the brain tissue, rendering e.g. surgical resection ineffective. While GBM cells possess large cell intrinsic migratory properties governed by cytoskeletal remodeling, adhesive bonds, etc., there might be additional driving factors determining migration, such as e.g. cell proliferation, pattern formation and cell density.

Past studies demonstrated that cells, including GBM, can undergo a transition from a fluid like, unjammed state with high cell mobility into an almost solid like, jammed state, blocking cellular mobility (2–11). There were multiple parameters proposed to induce jamming transitions, such as an increase or decrease of cell-adhesion, tension or increasing cell density (2–11). These studies suggested that increasing cell density regulates traction forces, thus leading to a migratory arrest for high cell densities (8, 11). Elucidating the role of cell density in migration is crucial, as (un-)controlled proliferation and thus increases in cell density is a universal process not only in development, but also in tumors. Yet, how cell density changes affect cellular migration is not completely understood (4, 7, 10, 12, 13).

The results of our previous studies indicate that cell density can lead to a cell-type dependent migratory arrest in GBM cells (10). Other research groups identified proliferation events as a factor associated with continuous fluidity of cell layers, and thus persistent cell mobility under confinement (14–18). When entering mitosis, cells detach from the extra-cellular matrix, round up, build the mitotic spindle and divide (19–21). The rounding process depends on a decrease in cell-cell and cell-substrate adhesion, and the parallel increase in intracellular pressure (22, 23), locally generating forces (24). In endothelial cells, division events were identified as sources of active stress generating vortex like flow patterns (15, 25) of sizes corresponding to several cell diameters (26). Notably, in some epithelial models it was argued that a decline in proliferation rate could lead to a glass transition, resulting in migratory arrest or that proliferation events are the main drivers of cellular reorganization (14, 24, 27).

The property to self-align is another factor potentially involved in the sustained migration of GBM cells under confinement. Notably, we observed previously that GBM cells show signs of self-organization, akin to “oncostreams” (28). Oncostreams are self-organized structures, emerging in glioma *in vivo* and *in vitro*, in which cells align parallel to each other and move in parallel or anti-parallel streams (29–31). Oncostreams were correlated with the grade and migratory capacity of gliomas (31). Interestingly, oncostreams in glioblastoma were later found to be reminiscent of a liquid-crystal like structure (32). Given the fact that GBM have a high cell density *in vivo*, such forms of self-organizations might promote an escape from the cell-density mediated migratory arrest. Indeed, this idea was supported by theoretical work and experimental studies showing cell density as a main inducer of oncostreams, but they can be abrogated by disturbing e.g. calcium signaling or actin-cytoskeletal organization (29, 30). Yet, the role of cell density, oncostreams and self-organization in GBM migration has only been addressed sparsely.

Here, we analyze the effect of different types of cell density fluctuations on the migration of GBM cells, including the effect of local, small-scale fluctuations, in the form of cell division events, cell density gradients and different absolute values of cell density. Furthermore, the relation between cell density, self-organization and escape from migratory arrest was assessed.

## Methods

### Cell culture

For experiments, LN229 and U138 GBM cells and primary GBM lines (GBM4, GBM10) were used. LN229 cells were purchased from the American Type Culture Collection (ATCC, CRL-261, Manassas, VA, USA) and U138 cells were obtained from Cell Lines Service (Cell Lines Service, 300363, Eppelheim, Germany). LN229 cells were cultured using 89% (v/v) Roswell Park Memorial Institute medium (Lonza, Basel, Switzerland, BE12-115F), supplemented with 10% (v/v) fetal bovine serum (FBS, Gibco, Carlsbad, CA, USA, 10500-064) and 1% (v/v) penicillin/streptomycin (P/S, Gibco, Carlsbad, CA, USA, 15140-122). U138, GBM4 and GBM10 were cultured in 89% (v/v) Dulbecco’s Modified Eagle Medium (Invitrogen, Waltham, MA, USA, 41965-062), and supplemented with 10% (v/v) FBS and 1% (v/v) P/S.

Primary GBM were derived from human biopsies as reported earlier (33). All patients provided written informed consent. The study was conducted in accordance with the Declaration of Helsinki and was approved by the local Ethics Committee of the Martin Luther University Halle-Wittenberg (project reference number: 2015-144).

For experiments involving mitomycin C treatment to inhibit proliferation, cells were subjected to a non-lethal dose of the substance, corresponding to 0.1 µg/ml and were continuously incubated and imaged with mitomycin C (34).

### Single Cell Migration

For measuring and analyzing single cell migration 1,000 cells were seeded per well in a 12-well plate (Greiner, Kremsmünster, Austria) 24 h prior to the start of experiments. Individual cells were imaged every 15 min with a microscope (Leica DMi8, Leica, Wetzlar, Germany) equipped with CO_2_ (5% (v/v)) and temperature (37 °C) regulation. The average cell speed and directionality were calculated based on cell tracking as described before (35). The directionality was defined as the ratio of the total distance travelled and the sum of incremental distances the cell moved between successive frames.

### Collective Cell Migration

For collective cell migration experiments 450,000 cells were seeded per well in a 12-well plate. Twenty-four hours afterwards cells were imaged with an inverted microscope (DMi 8, Leica, Wetzlar, Germany), in a fully humidified atmosphere with 5% CO_2_ (v/v) and at 37°C. Images were acquired every 3 min for 60 h. For analyzing cell migration particle image velocimetry, based on PIVlab, was used with a final cross-correlation window size of 16 × 16 pixels (pixel size: 0.48 µm), as described elsewhere to obtain local velocity fields (10, 28, 36).

To assess the effective motion of cells the mean squared displacement (MSD) 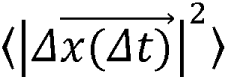 and its scaling coefficient α(Δ*t)* were evaluated, using the following equation:

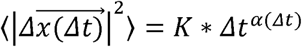

With the generalized diffusion coefficient *K*. The scaling coefficient retrieves information about the type of motion observed inside the monolayer, with sub-diffusive motion for α*<1*, diffusive motion for α≈*1* and super-diffuse motion for α*>1*.

### Identifying Proliferation Events and Cell Density Dependent Migratory Properties

To automatically identify individual proliferation events, a machine learning model was used as described previously (28). Briefly, individual proliferation events in a dense monolayer were identified using a three-step process, consisting of an initial u-Net based segmentation, to reduce the region of interest and speed up evaluation, followed by candidate extraction using 2D cross-correlation. Finally, candidate divisions were classified as true or false positives using a GoogLeNet classifier. To match division events occurring over multiple frames the Munkres global nearest neighbor algorithm was used for detection to track assignment. As cell divisions can be incomplete or cells can be excluded from the layer having a similar appearance as mitotically rounded cells, but with a significantly longer visibility, detections that were present for more than 60 min were discarded.

To assess the influence of individual proliferation events on the layer velocity, for each identified proliferation event the speed around the division was calculated as a function of distance to the division and normalized to the speed of the rest of the layer. The same procedure was applied to up to 20 images, corresponding to 60 min, before and after the detection of the mitosis event, corresponding to the assessment of the effect of pre-mitotic contraction and post-mitotic expansion. To further assess the cell density in each field of view, the initial cell number at t = 0h was manually counted in a sub-field of 144×144µm² and extrapolated to the whole field of view. To calculate the cell density ρ over the whole time for each field of view the number of proliferations in each field of view, normalized to the surface area was added to the initial cell density. Combined with the layer migration speed *v* this approach allowed to calculate the scaling coefficient α between both quantities as follows:

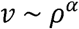

### Analyzing Self-Organization in Monolayers

To determine local orientation of cells in a layer the largest eigenvector of the structure tensor *J* of the image *I* was used. The structure tensor *J* was defined as (37):

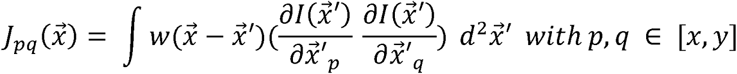

and the Gaussian window function *w*. From this cellular orientation, the (local) nematic order parameter *S* was calculated as follows:

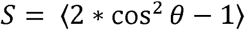

With θ being the angle between the cellular orientation and the mean orientation of all cells, either in the whole field of view or a defined sub-window. To evaluate the spatial dependence of the order parameter, a power law was fitted as *S(l)∼l^a^*, with the decay exponent *a* and the length scale *l* (32). For a long range order *a* approaches 0 and *S* remains constant. For 0>a>-1 the order is considered to be of a quasi-long range type, with the extreme case a =-1 corresponding to a random order of orientations. For short-range order, an exponential decay is expected, but for the data given here, a power law describes the data best.

As nematics contain topological defects, ±1/2 defects were identified using the Matlab toolbox “defector find” (38) on the previously calculated cellular orientation fields. This toolbox calculates the winding number around potential topological defects in order to classify them as ±1/2 defects. Notably, this allows assessing evolution of the number of topological defects as a function of time and the movement of topological defects.

To evaluate if the ordering might be of a polar type, a polar order parameter *V* was introduced (29):

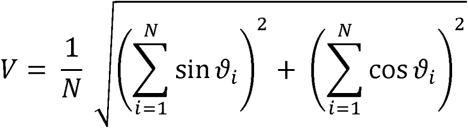

Here, *N* corresponds to the number of vectors □ in the whole field of view or a defined sub-window. □ is the angle between cellular orientation and velocity at the same spatial position. Similar to the nematic order parameter, the decay of the polar order parameter as a function of coarse graining was assessed. Here, an exponential decay function fitted the data best:

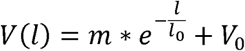

With the characteristic decay length *l_0_* and the constant offset *V_0_*.

### Cell Exclusion Assay

For cell exclusion assay 50,000 cells were seeded per well in a confined ring structure in 12-well plates. Twenty-four hours after cell seeding, the cells reached a confluent status. Thereafter, the ring structure was removed, and cells were transferred to the microscope (Leica DMi8, Leica, Wetzlar, Germany, 5% CO_2_ (v/v), 37 °C) for monitoring cell expansion. Five regions of interest were imaged per well. Cell expansion was monitored for 60 h with images taken every 5 min.

To analyze cell expansion an u-Net was trained to segment the cell layer from the background. Local velocities and other quantities were calculated as described under “Collective Cell Migration”, with the restriction of the analyzed area to the cell covered area.

For assessment of cell density dependent effects another u-Net was trained to identify cell-cell boundaries in phase contrast images and thus assess cell size inside the cell layer, based on 52 manually segmented sample images. To further improve the detection of cells, too small (less than 120µm²) and too large (more than mean size + 2 * median absolute deviation of size) were discarded. Furthermore, cells have been assigned to tracks via the Munkres global nearest neighbor algorithm and cells that have not been assigned in at least 4 of the 5 following images were discarded too, to reduce the number of misdetections. The following energy function was used as input for the Munkres algorithm for the assignment of a detection *d* to track *t*:

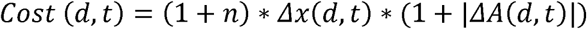

Here, *n* corresponds to the number of frames the detection *d* was not assigned to any track *t*, Δ*x(d,t)* the normalized distance of the detection d to the last detection of track *t* and Δ*A(d,t)* corresponds to the normalized difference in cell size between the currently detected cell *d* and the last detection of track *t*. If the assignment cost for a detection was too high, a new track was created.

To calculate the gradient in cell density in a field of view, the cell density in dependence of the distance to the cell cluster boundary was calculated and linearly fitted. The slope of the fit was used as cell density gradient estimate.

## Statistics

Statistics were performed using the two-tailed ANOVA with the Tukey post-hoc test or the two-sided sign test. Significance was defined for p < 0.05. All error-bars and shaded areas depict the standard error of the mean. Experiments were repeated at least three independent times. For live-cell imaging experiments assessing collective motion, 5–8 fields of views were measured per experiment and condition. Confidence intervals for fit parameters were obtained using the MatLab function confint.

## Results

### Glioblastoma Show Cell Type Dependent Response to Increasing Cell Density

To establish the migratory potential of all four GBM cell lines, we first analyzed the single cell motility. GBM4 and U138 moved significantly faster and their movement pattern tended to be more directed than that of LN229 and GBM10 (Fig. 1A, Fig. S1). While the overall movement speed in a dense monolayer does not follow the same pattern, with GBM10 being the fastest and LN229 cells slowest, the overall movement for GBM4 and U138 cells is more effective, as demonstrated by the higher MSD and scaling coefficient (Fig. 1B-E). Thus, the overall notion seemed to be that U138 and GBM4 move significantly more effective than LN229 and GBM10. Of note, while the overall speed of GBM cells seems to mostly decrease over time and thus cell density, when analyzing the scaling coefficient of the mean squared displacement at the beginning and end of the measurement period, corresponding to low and high cell densities, a cell type dependence was found. Both LN229 and GBM10 showed lower scaling coefficients over time, approaching a diffusive type of motion for high cell densities, while U138 and GBM4 cells were mostly unaffected in their movement pattern (Fig. 1F-I).

**Figure 1:**
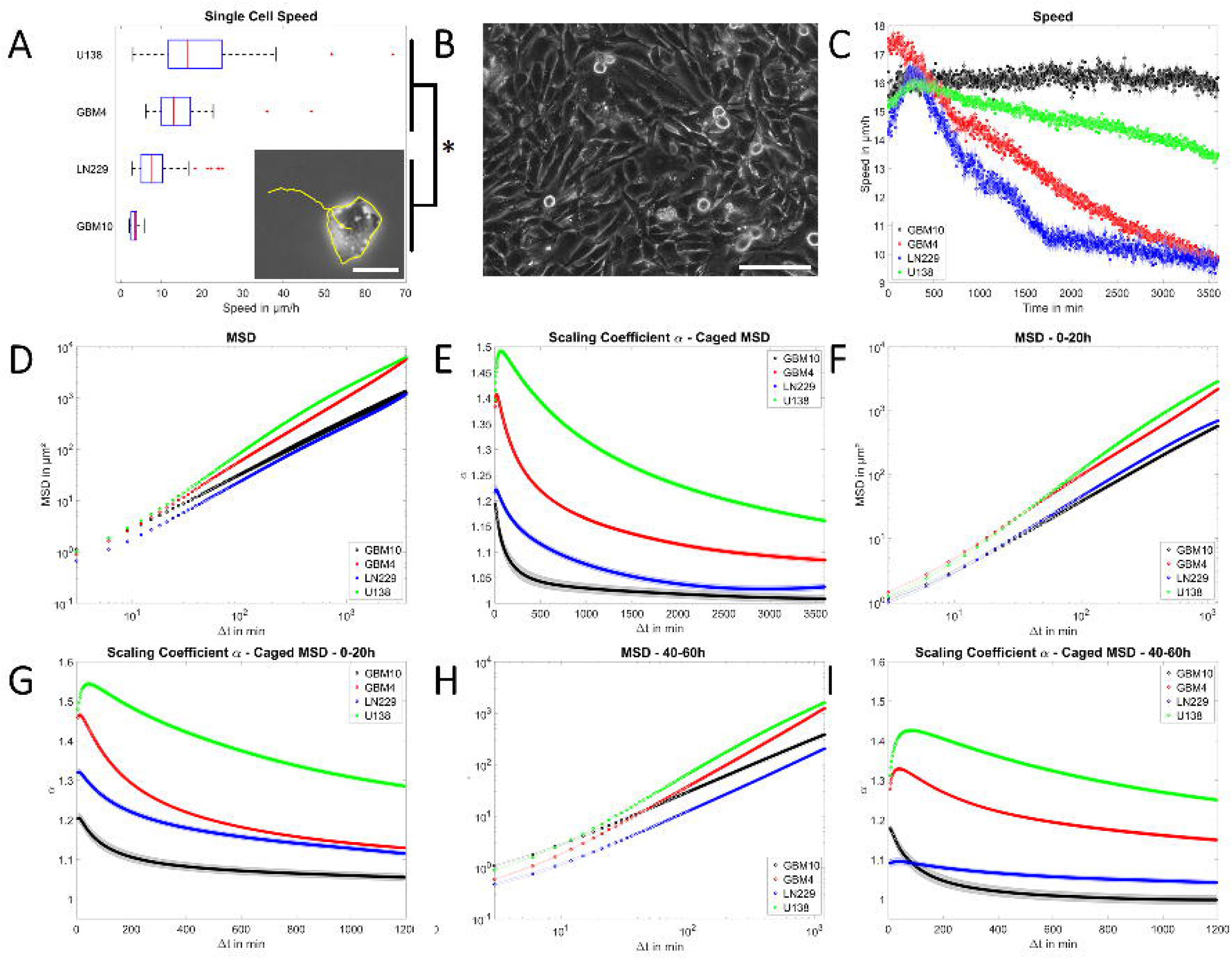
Migratory capacity of GBM cells. A) Graph of the single cell migration speed, with the inlet showing the path and edge detection of a LN229 cell. U138 and GBM4 cells are significantly faster than LN229 and GBM10. Scale bar corresponds to 25 µm. B) Image of a dense monolayer of U138 cells at 0 h, illustrating the experimental starting point. Scale bar corresponds to 125 µm. C) Depiction of the mean layer speed as a function of time for all 4 GBM cell lines. D) and E) Analysis of the mean squared displacement and the scaling coefficient of all cell lines for the whole 60 h time frame. F) to I) Illustrations of the same parameters as in D) and E) but for the first 20 h for F) and G), or the time from 40 h to 60 h in H) and I). Please denote that the movement pattern of U138 and GBM4 cells is less affected than that of LN229 and GBM10 by increasing time and thus cell density. Error bars and shaded areas depict the standard error of the mean. Box plots show the median (red line), 25 and 75 percentile (box), non-outlier range (whiskers) and outliers (red dots). Sample sizes: A) n_U138_ = 44; n_GBM4_ = 47; n_LN229_ = 83; n_GBM10_ = 43; C) to I): n_U138_ = 184; n_GBM4_ = 139; n_LN229_ = 79; n_GBM10_ = 60;

Consequently, we next evaluated different aspects of a changing cell density to elucidate a potential course for the cell type dependent response. Therefore, three different types of changes in cell densities were assessed: 1) Local changes of cell density caused by individual proliferation events, 2) response of cell layers when subjected to cell density gradients in a cell exclusion assay and 3) the effect of a global increase in cell density caused by the accumulation of all proliferation events.

### Proliferation Events Cause a Local Acceleration in Glioblastoma

As proliferation is an ubiquitous process present throughout all our experiment we evaluated first if isolated proliferation events affect the migration of GBM cells. To test this hypothesis, a dense, homogenous monolayer was analyzed, which is largely independent on cell density gradients. For this analysis a data set consisting of 462 regions of interest, including approximately 134,000 cell division events were analyzed (Table S1).

To analyze the effect of proliferation events, we first validated our approach to detect division events. Therefore, we manually determined the cell density at t = 0h for each region of interest (Fig. S2), and extrapolated the cell density for all subsequent measurement times using the described approach to detect cell division events. This allowed us to assess the number of proliferation events and calculating and comparing the scaling coefficients between cell density and layer speed for U138 and LN229 cells, with those previously measured and published (10) (Fig. 2A,B). Scaling coefficients differed by only ±0.01, being in very good agreement with the previously made manual estimates. Thus, in addition to previously made tests (28), this result showed the capability of our system to track division events reliably. As can be further seen by the results, GBM10 and LN229 cells are significantly more proliferative than GBM4 and U138, with GBM4 even showing signs of a growth arrest (Fig. 2A).

**Figure 2:**
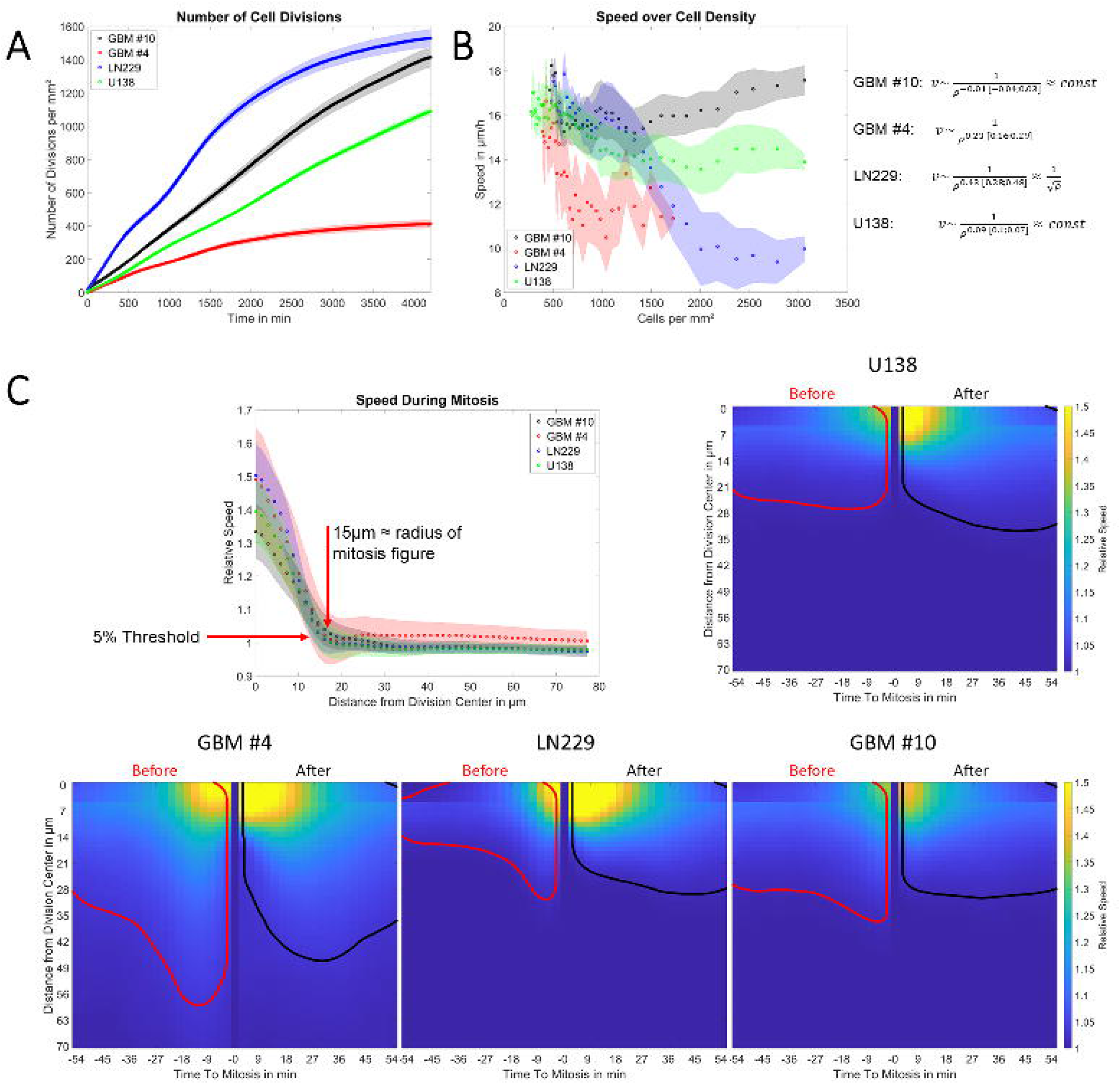
Analysis of proliferation and its effect on migration speed. A) Graph of the number of proliferation events as a function of time. Please denote that the less-migratory LN229 and GBM10 cells have a higher proliferation rate. B) Plot of the layer speed as the function of cell density, showing cell type dependent behavior and validating previously found scaling coefficients. C) Quantification of the effect of proliferation on velocity fields. The top left graph demonstrates that in the state of being mitotically rounded those cells do not significantly affect the motility of cells around them. The remaining graphs illustrate the effect of pre-mitotic cell contraction and post-division spreading. The encircled regions depict the spatial and temporal regime were pre-mitotic contraction (red) or post-mitotic spreading (black) led to a measurable increase in velocity (>5%) in the surrounding cells. Error bars and shaded areas depict the standard error of the mean. Sample sizes: A) to B): n_U138_ = 184; n_GBM4_ = 139; n_LN229_ = 79; n_GBM10_ = 60; C) n_U138_ = 58,131; n_GBM4_ = 16,645; n_LN229_ = 34,993; n_GBM10_ = 24,612;

To assess qualitatively if proliferation events could be associated with changes in the migratory properties of cells a first visual examination of the velocity field of cells around proliferation events was done (Fig. 3). Three distinct phases of relevance were identified: a phase of pre-mitotic contraction, a mitotically rounded phase, hereafter termed mitosis event, and a post-mitotic expansion phase. During the pre-mitotic contraction, the velocity of the surrounding cells increased (Fig. 3, marked red), but when cells were fully mitotically rounded the velocity slowed down again (Fig. 3, marked dark green) and during daughter cell expansion an increase in the speed of the surrounding cells was observed (Fig. 3, light green) before an eventual normalization. These qualitative observations motivated the distinction of the three different phases mentioned above for the quantitative analysis below.

**Figure 3:**
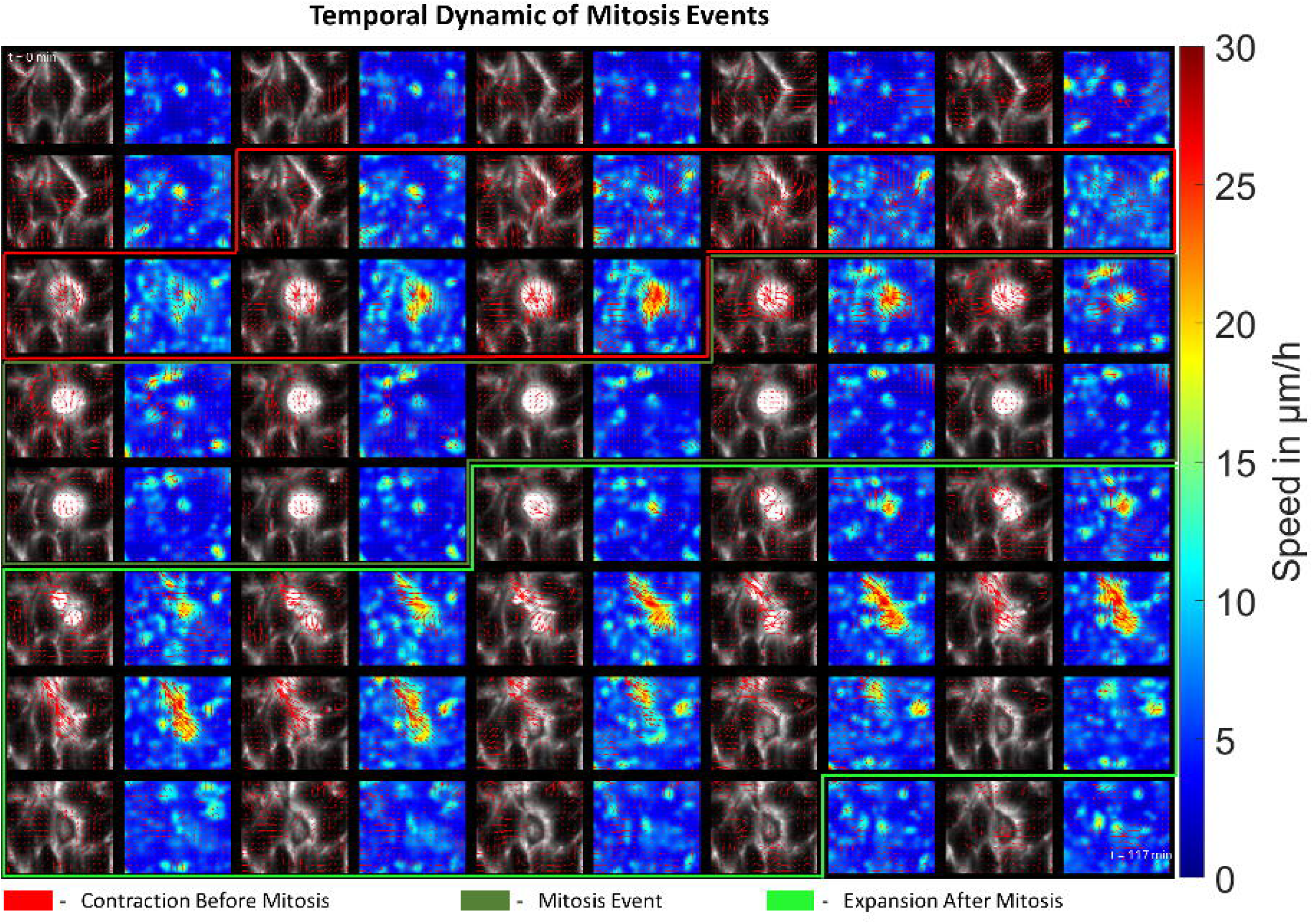
Temporal Dynamic of Mitosis Event. This graph illustrates the temporal dynamics of a mitosis event over 2 h, combined with the accompanying changes in the surrounding velocity field. It can be found that the process leading to mitotic rounding affects the speed of the surrounding cells, as does the daughter cell expansion after mitosis.

For quantification of the proliferation-induced changes of the velocity field, the speed was analyzed as a function of the distance to the dividing cell during mitosis, which is the time a cell was identified as mitotically rounded, for all regions of interests and all cell densities. The speed around the mitosis event was found to be significantly higher at the core of the event, but dropped quickly, reaching a value that is mostly indistinguishable (≤5% increase) from the background speed for distances of 15µm or higher, corresponding roughly to the radius of the mitosis figure (Fig. 2C). Thus, during mitosis the overall layer migration speed was not altered significantly. Nonetheless, based on these measurements the drop off point to 5% appeared to be a reasonable cut-off level for determining a region that is affected by proliferation events.

Afterwards the time up to 1 h before the start and up to 1 h after the end of mitosis was analyzed and the distance dependent speed calculated. There, both pre-mitotic contraction and post-mitotic expansion led to an increase in the speed of the surrounding layer, affecting multiple cells around the dividing cells (Fig. 2C). Of note, those effects can begin longer than 1 h before the start or after the end of the mitosis event. Interestingly, the spatio-temporal pattern of both parts was different. The effect of pre-mitotic contraction tended to be smaller the further in time the mitosis event was and reached its peak, both in effective range and overall magnitude, approximately 10 min prior to mitosis. The effect of the post-mitotic expansion on the other hand built up its effective range over time, reaching its maximal range approximately 30 to 40 min after mitosis (Fig. 2C).

To elucidate whether the effects of cell divisions are dependent on the global cell density, we analyzed the maximal effect range, that is the highest distance with an increase in speed ≥5% relative to the background, as a function of cell density. It was found for all four cell lines and pre-and post-mitotic phase that increasing cell density reduced the range in µm affected by a division event. Yet, this effect was counterbalanced by the increasing cell density, yielding an effective increase in the number of cells being affected by a nearby proliferation event. Thus, cells three or more cell radii away from a division event could be influenced by it (Fig. S3).

Having obtained the average temporal and spatial range of a division events, we calculated the average area affected by proliferation events for each individual field of view. Approximately 20-30% of the monolayer were on average affected by cell division events for each cell line (Fig. S4). Notably, these numbers were used – together with the data from Fig. 2C – to estimate the proportion division events had on the total speed. The overall proportion of cell division events turned out to be ≈4-5% for all cell lines, suggesting that individual proliferation events were no main driver of cell migration in GBM in our system and cannot explain the phenotypic differences between the cell lines. Thus, we next analyzed the effect of cell density gradients in cell migration.

### Cell Density Gradients Can be a Driver of Glioblastoma Migration but in a Cell Type Dependent Manner

To generate a cell density gradient, cells were seeded in a confined circular space and after removal of the confinement the expansion of the layer was measured. Analysis of the cellular speed as a function of distance to the cell front showed that the layer expansion was initiated at the front and gradually spreading deeper into the cell layer (Fig. S5). In the presence of proliferation, it was found that the previous hierarchy in terms of migration was no longer preserved, showing that U138 and LN229 cells were expanding faster than GBM4 and GBM10. Given the fact that LN229 and GBM10 had the highest proliferation rate, combined with lower migratory capacity compared to the other cell lines it indicates that cell density gradients at or near the cell front could drive layer expansion. To test this hypothesis proliferation was inhibited by addition of a non-lethal dose of 0.1µg/ml mitomycin C, as verified by visual inspection of the images. Initially, mitomycin C treated cells behaved similar to the control cells, up to ≈2000-2200 min. Then, for all cell lines but U138 the mitomycin C treated cells were expanding significantly slower than control cells, and for LN229 and GBM10 an arrest of expansion was observed (Fig. 4A). Notably, in the absence of proliferation U138 and GBM4 were expanding faster than LN229 and GBM#10, restoring the previous notion in terms of migratory capacity. To demonstrate the dependence of the migratory phenotype on the actual cell density gradient at or near the cell layer boundary an u-Net to segment individual cells inside of the monolayer was trained. Testing the neuronal network in an independent image batch, not used for training, an accuracy of 84.7% was found. Together with the sized based filtering and track assignment, the detection gave good results for all but one cell line (Fig. 4B). Only for GBM4, no good segmentation results were obtained, as no manual ground truth could be created, as cell edges were often not clearly discernable (Vid. S1).

**Figure 4:**
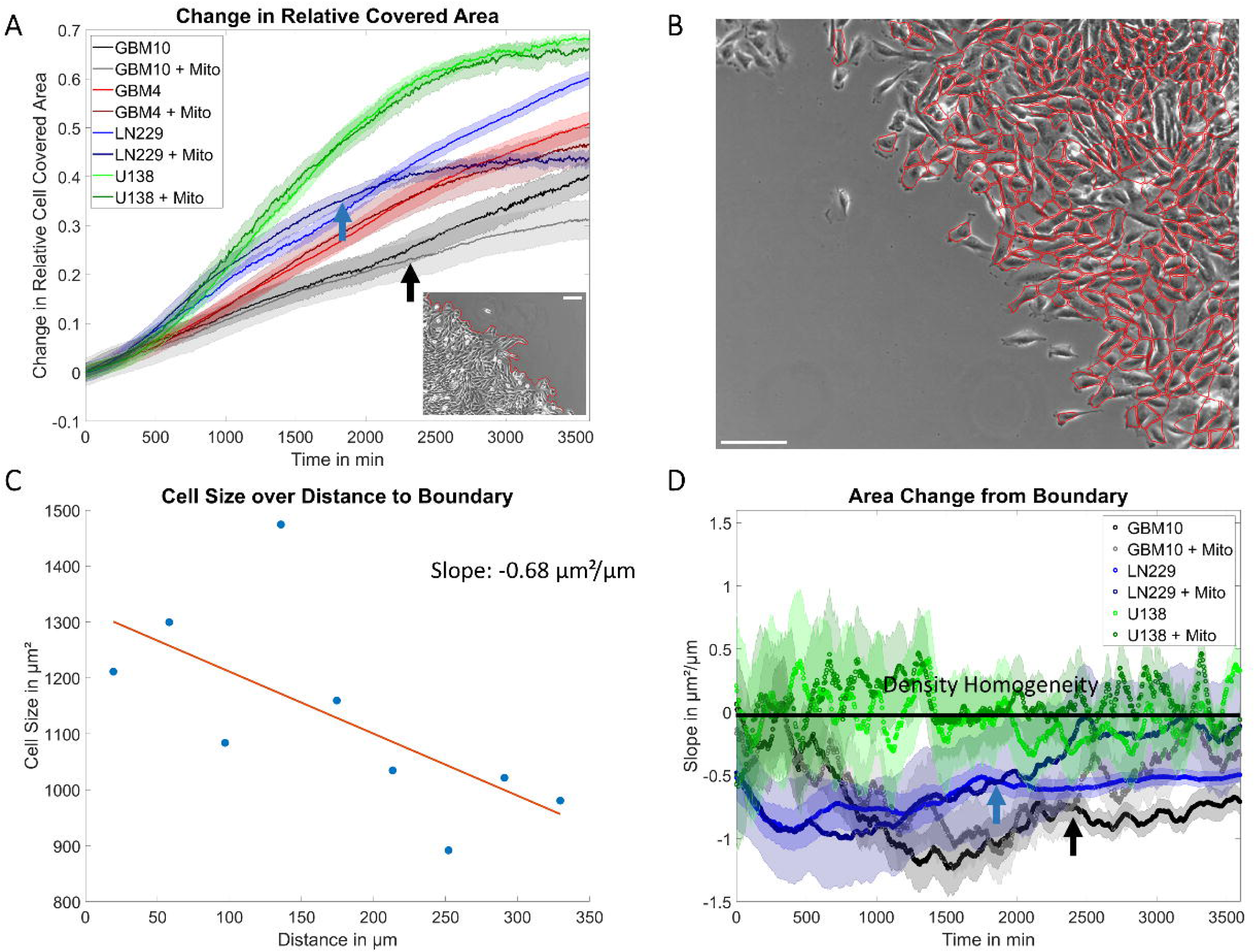
Analysis of GBM cell expansion in presence and absence of proliferation. A) Shows the expansion rate of GBM cells under control conditions and when treated with mitomycin C to block proliferation. For U138 saturation occurs as the whole field of view is eventually covered by the cells. Blue and Black arrows show the same time point as in D), roughly corresponding to the points were the expansion rate of mitomycin C treated LN229 and GBM10 cells diverged from the controls. The inlet shows a typical sample image of the cell exclusion assay for LN229 and the associated detected cell front. Scale bar corresponds to 100 µm. B) Image of the final cell segmentation for LN229 cells, showing good segmentation results. Scale bar corresponds to 75 µm. C) Plot of the cell size as a function of distance of the cell to the cell front for LN229 cells, including a linear fit and its slope, corresponding to the cell density gradient. Please denote the decline in cell size with increasing distance from the front. D) Graph of the cell density gradient as a function of time for mitomycin C treated and untreated cells. The black line shows the reference for the absence of any cell density gradient. Arrows mark the time points at which the cell density gradients between the control and mitomycin C treated LN229 and GBM10 cells start to diverge. Error bars and shaded areas depict the standard error of the mean. Sample Sizes: A), D) n_U138 CTL_ = 24; n_U138 Mito_ = 11; n_GBM4 CTL_ = 39; n_GBM4 Mito_ = 10; n_LN229 CTL_ = 29; n_LN229 Mito_ = 15; n_GBM10 CTL_ = 13; n_GBM10Mito_ = 13;

Based on this segmentation, we calculated the cell size as a function of its distance to the cell cluster edge for each image and subsequently calculated its slope, as a measure for the cell density gradient (Fig. 3C). For LN229 and GBM#10 cells in control conditions, the cell density gradient decreased after removing the confinement, reaching a minimal value and afterwards increasing to a constant level at roughly of-0.5µm²/µm or-0.7µm²/µm, respectively (Fig. 4C). Initially, mitomycin C treated cells behaved similarly, until approximately 1850 min or 2300 min after treatment, when the cell density gradient started to increase to higher values than the previous equilibrium value, eventually reaching a state of homogeneous cell density. Notably, the time the cell density between control and mitomycin C treated cells diverges corresponded to the time point the cluster expansion diverged as well. Upon reaching the equilibrium cell size, cluster expansion came to a halt, suggesting that expansion of GBM10 and LN229 was mostly driven by the cell density gradient (Fig. 4C). This was also supported by the increase in average size for both cell lines, after mitomycin C treatment and the scatter plots of the cell density gradient over the expansion rate (Fig. S6), showing a negative correlation. While, GBM10 and LN229 cells behaved qualitatively similar, for U138 cells, no significant deviation of the cell density gradient from zero was observed over time, independent on the presence or absence of proliferation. Thus, the expansion of this cell line is not determined by the cell density gradient (Fig. S6B) present throughout the cell exclusion assay. Consequently, this suggests that the outward expansion of U138 cells was mostly driven by active migratory processes while for LN229 and GBM10 expansion was governed by the passive pressure of the growing cellular mass. Notably, no large qualitative differences were found in cluster cohesiveness between groups, capable of explaining those differences. For all cell lines, a high cohesiveness of the front was observed, with only few cells escaping the layer (Vid. S1-S5).

These experiments suggested that the cumulated effect of individual mitosis events, leading to increased cell density and here to cell density gradients, could be strongly involved in cell migration.

### Glioblastoma Show Self-Organization in Liquid Crystal-Like Structures

To analyze the effect of increasing cell density, independent of cell density gradients, we turned back to the confluent monolayer system of homogeneous cell density. As observed in the cell exclusion assay (Vid. S1, Vid. S2) and in a previous study of ours (28), both GBM4 and U138 showed signs of self-organization for high cell density, reminiscent of oncostreams (31), not visible in the other two cell lines (Fig. S7). Thus, we reasoned those structures might be involved in the escape from cell-density induced migratory arrest. To assess the type of structures formed by GBM4 and U138 cells the monolayers were analyzed manually to identify breakage points in the underlying structures, termed topological defects. Three types of structural defects were identified: +1/2 defects (comet),-1/2 defects (trefoil) and +1 defects (aster, Fig. 5). From those, the vast majority were of the ±1/2 type and therefore we reasoned the underlying organization to be of a nematic type. Consequently, the nematic order parameter was used to assess the evolution of structure formation over time in our monolayer models. Indeed, in GBM4 and U138 the nematic order parameter increased over time and thus cell density, regarding the whole field of view (≈0.29 mm², Vid. S6, Vid. S7). The other two cell lines showed only little nematic ordering that was slightly increasing over time (Fig. 6A, Vid. S8, Vid. S9). Analyzing the nematic order for different spatial dimensions ranging from windows with edge length of 25 µm to 325 µm a decreasing nematic order was found when decreasing window size. For LN229 there was a significant increase in the local nematic order over time, spanning cell populations of the size of up to ≈0.04 mm², implying the evolution of a locally ordered state (Fig. 6B). For GBM#10 the nematic order did not change strongly over time, even when regarding small windows (Fig. 6B). To evaluate the range of order further the decay of the order parameter in dependence of the analyzed block-size was calculated. This decay function was best fitted by a power-law, with values of the exponents approaching ≈-0.1 for GBM4 and U138, implying a high order on a large scale (Fig. 6C). For LN229 cells, the scaling coefficient increases from-0.5 to-0.35, implying a gradually stronger ordering (Fig. 6C). In contrast, in GBM10 the scaling coefficient stayed roughly constant around ≈-0.55 for all times investigated (Fig. 6C). These findings are in line with the qualitative assessment of the emergence of order from before. To further validate the increasing organization of GBM cells over time, the number of ±1/2 defects as a function of time was evaluated. This demonstrated a strong and steady decrease of defects in U138 and GBM4 over time, a less pronounced decrease in LN229 and a constant number for GBM10, being in line with the expectations based on the nematic order parameter (Fig. S8-11E). Of note, the number of +1/2 and-1/2 defects was, as expected, almost identical for all cell types validating these results further. To assess if the defects themselves were drivers or inhibitors of motion, the MSD and movement path of defects were analyzed relative to the defect orientation for defects persisting at least for 3 h. Defects showed a sub-diffusive movement pattern, with no favored direction relative to their orientation (Fig. S8-11A-D).

**Figure 5:**
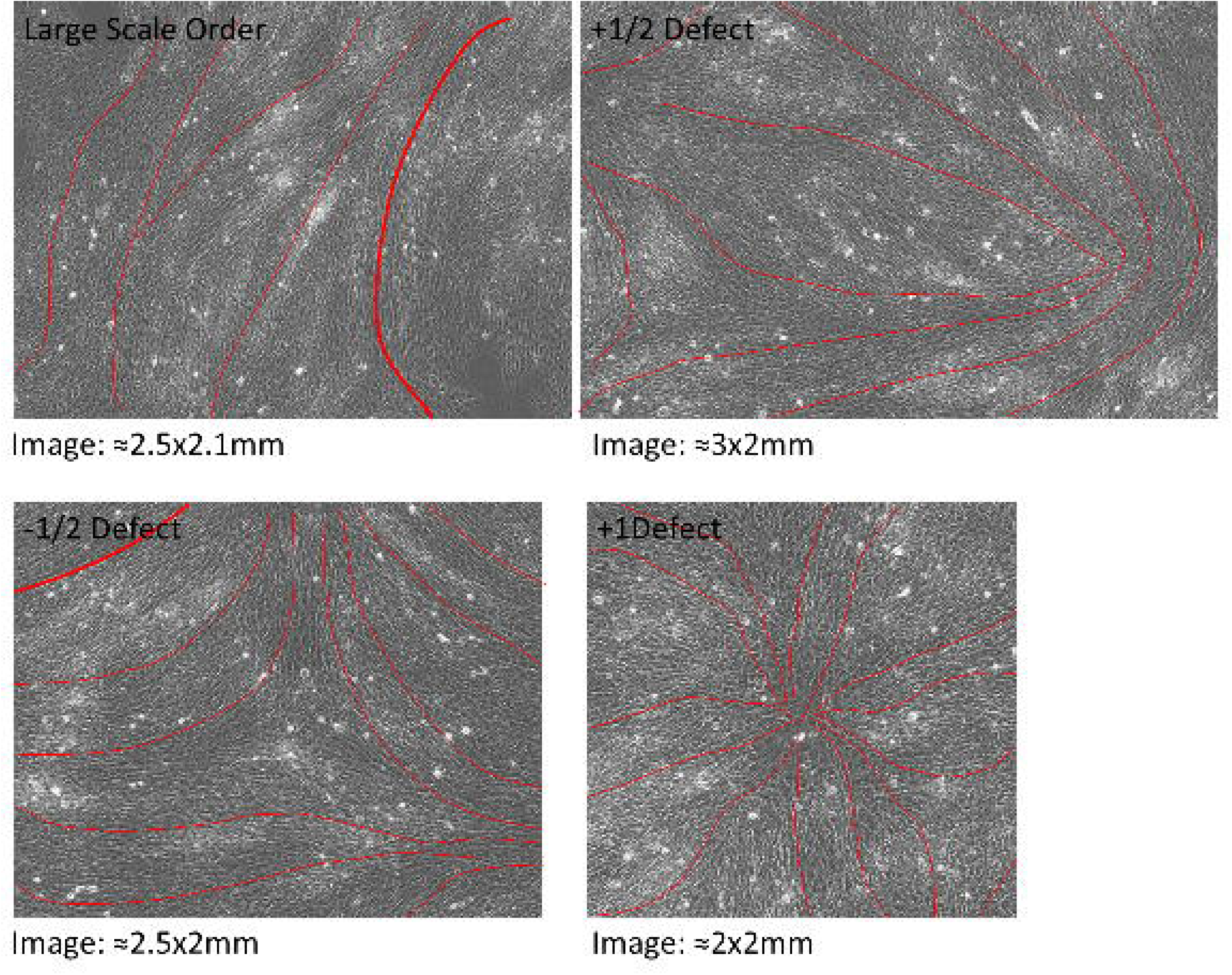
Topological defects found in GBM cultures. The top left image shows GBM4 cells in a highly organized state, mostly aligned in one direction. The top right image depicts GBM4 cells organized in the form of a +1/2 or comet like defect. The bottom left image illustrates the formation of-1/2 or trefoil defects in GBM4 cells, and the bottom right image is that of a rarely observed +1 or aster defect.

**Figure 6:**
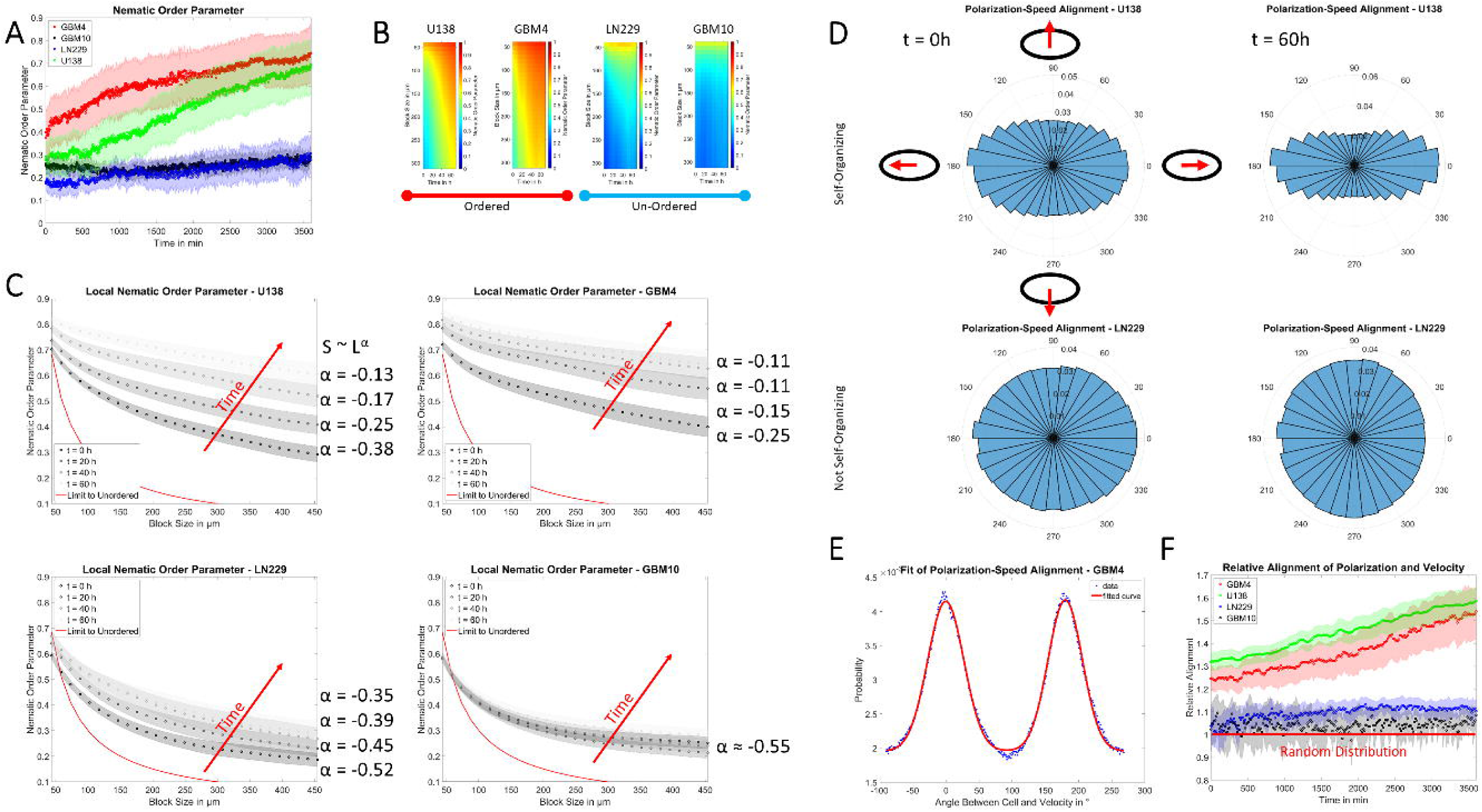
Nematic order in GBM cells. A) Plot of the temporal evolution of the nematic order parameter for the whole field of view. B) Heatmaps of the temporal and spatial evolution of the order parameter for all four GBM lines. C) Graphs of the nematic order parameter as a function of the used spatial scale and defined time points. Red lines show the expectation value for a random distribution of orientations and the red arrow depicts the direction of increasing time. To the right of the graphs the exponents of the power law fits for each chosen time are shown. D) Polar plots of the alignment of cellular orientations with the velocity vectors at the beginning (left) and after 60 h (right) for U138 (top) and LN229 (bottom) cells. E) Sample distribution of the angle between cell orientation and velocity for one field of view of GBM4 cells, together with the respective fit in red. F) Relative alignment of cellular orientation and velocity field, in terms of the number of cells moving in ±22.5° of their orientation, over time. The red line shows the value expected for a random alignment. Error bars and shaded areas depict the standard deviation. Sample sizes: A) to C), F): n_U138_ = 184; n_GBM4_ = 139; n_LN229_ = 79; n_GBM10_ = 60;

### Liquid Crystal-Like Self-Organization Promotes Migration in High Cell Density Limit

To test if the nematic order of GBM cells is related to the directionality of cellular migration, the alignment of velocity vectors and cellular orientation was analyzed. For LN229 and GBM10 the alignment of cells and velocity vectors resemble almost a random distribution. For GBM4 and U138 both vectors tend to align in a parallel and anti-parallel manner, with increasing alignment over time and cell density, implying the formation of anti-parallel moving streams in the monolayer (Fig. 6D). To quantify this phenomenon further the probability density function was fitted as the sum of two Gaussians and a constant offset (Fig. 6E). To quantify the proportion of cells moving in the direction of cellular alignment, the proportion of cells situated at the peak centers ±22.5° was calculated and normalized to the expectation value for a uniform distribution. For GBM4 and U138 the amount of cells moving in alignment with their orientation was significantly higher than expected by chance and increasing over time and thus cell density. For GBM10 this value was close to the value expected by chance, while a slight increase over time was observed for LN229 cells (Fig. 6F). As these measurements suggesting the emergence of anti-parallel moving streams of cells, it was evaluated if additionally a stronger polar order is emerging. Therefore, the polar order parameter for the velocity field was calculated. There, only small differences were observed between cell lines and the polar order was generally smaller than the nematic order, with no significant changes over time and thus cell density (Fig. S12). The decay of the velocity order parameter was best fitted by an exponential decay, implying only a short-range order and decay coefficients stayed constant for all cell types over time and had values between ≈90 and ≈140µm, depending on the cell line. Therefore, the emerging order in GBM cultures is mostly of a nematic and not of a polar type.

Thus, these measurements suggest that the emerging nematic order is related to a consistent movement of cells. If this is valid, it is expected that even inside a monolayer of one cell type, groups of cells showing high nematic order (S>0.7) migrate more effective than cell groups of lower order (S<0.3), albeit the effect is expected to be smaller than the difference between highly and lowly ordered cell lines (29). To test this hypothesis, all 462 regions of interest of size 0.29 mm² were tested if they contained regions of low and high nematic order at the same time (Fig. 7A). The regional threshold was set to 1820 µm² to contain multiple cells. The movement of those clusters was than tracked for the following 6 h. Interestingly, the highly ordered cells in U138 and GBM4 were slower (Fig. S13), but moved more effective, as shown by the higher mean squared displacement. The effectiveness of motion also was increasing over time and thus cell density (Fig. 7B). Similarly, ordered clusters of LN229 moved more effective, but no significant changes in speed were observed (Fig. 7B, Fig S13). For GBM#10 no such effect was observed (Fig. 7B), but due to the overall low ordering of this cell line only very few regions of interests containing both regions of high and low ordered cell groups remained (at lowest 21 of 60).

**Figure 7:**
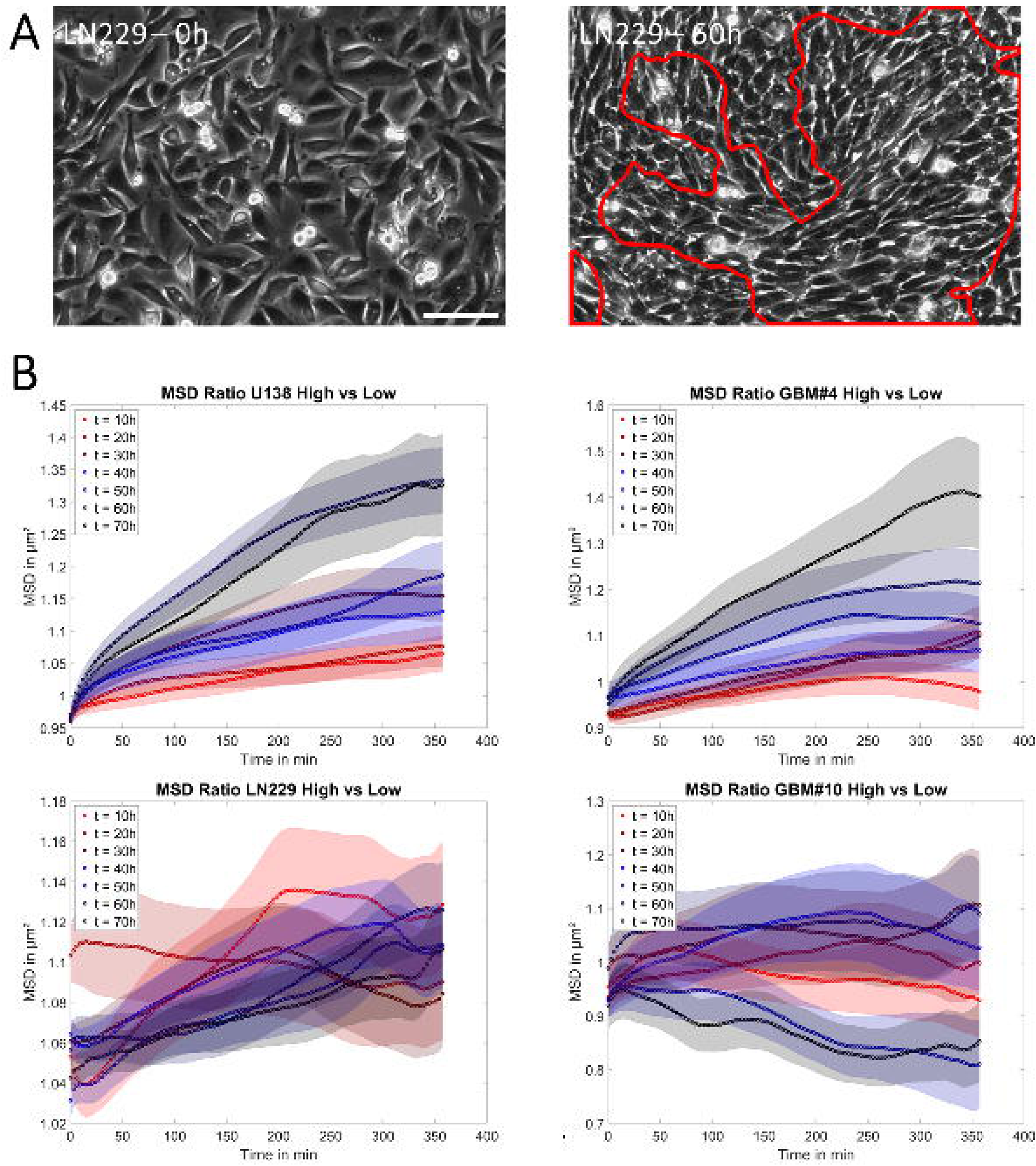
Comparison of the migration of ordered and unordered regions in one field of view. A) Example image of the emergence of local nematic order in LN229 cells. The left image shows the cell layer at 0 h, while in the right image taken after 60 h a locally ordered region, encircled in red, has developed. Scale bar corresponds to 100 µm. B) Ratios of the mean squared displacement for regions with high nematic order over those with low nematic order for U138 (top left), GBM4 (top right), LN229 (bottom left) and GBM10 (bottom right). Error bars and shaded areas depict the standard error of the mean. Sample sizes for t = 0,10,20,30,40,50,60h: B): n_U138_ = 180, 167, 161, 145, 135, 101; n_GBM4_ = 139, 131, 122, 115, 106, 88; n_LN229_ = 79, 79, 78, 78, 77, 73; n_GBM10_ = 32, 27, 28, 29, 28, 21;

## Discussion

In this manuscript, we analyzed the effect of various changes in cell density on the migratory capacity of GBM cells. Thereby, two different phenotypes were identified, with one being mostly independent on cell density, while the other one showed an effective slowdown of cell body translocation with increasing cell density. Our analysis suggested that organization of cells in a liquid crystal-like manner prevented the cell density induced migratory slow down.

Under confinement and with increasing cell density, several studies suggested that cellular movement slows down, eventually leading to a migratory arrest (4–11). Yet, other studies showed opposing results with cells staying motile even for high cell densities (10, 39).

Nonetheless, for cells to be able to escape the confinement of their neighbors for high cell densities, sufficient forces need to be generated. One potential course for such fluctuations are proliferation events (27). Thereby, cells were considered to generate forces mostly in different phases. Starting with the process of mitotic rounding leading to a net inward stress (17, 18, 26), caused by acto-myosin contractility and increased hydrostatic pressure (21, 40–44). After division, during spreading and reintegration of the daughter cells into the layer, cells generate outward stress (17, 26). Notably, for both phases we did observe an increase in the local velocity around the dividing cell, supporting this notion and validating our approach. The tension around a dividing cell was found to change by around 10-20% in a distance of up to two cell diameters from the division event, being very similar to our estimated effect size of 2-3 cell sizes (45). Past studies demonstrated that in certain epithelial cells and during development, proliferation events are sufficient and/or necessary for the induction of cellular reorganization and thus a migratory phenotype (14, 24, 27). Here, we could confirm an effect of proliferation events on the local velocity field, spanning a similar spatial scale as in MDCK cells (15), albeit here the effect was very small on the tissue scale. A potential reason for this discrepancy might lie in the used cellular system. For this study, we used highly motile GBM cells, moving at speeds between 10 to 18 µm/h, while in the aforementioned studies healthy cells with lower migratory capacity were used. E.g. in a very similar setup MDCK cells moved at most ≈6 µm/h (27), implying a significantly lower cell-intrinsic force generation capacity, helping to overcome cell density induced confinement. To the authors’ knowledge, there is only one study on mamma carcinoma cells evaluating the impact of proliferation events on migratory properties of tumor cells (46). In their study, West et al. reported similar speeds for the monolayer as observed here, and found the effect of proliferation events to drop off in a distance of around one cell diameter, with a similar magnitude of the effect of at most ≈50-70% increase in speed. Albeit similar, the effective range was slightly lower, potentially due to different biological behavior of the used tumor entities or the significantly lower number of analyzed division events. Thus, all results obtained here are in good agreement with past studies, showing a local effect of proliferation events on the velocity field, but here cell division did not appear to be a main driver of migration, likely due to the higher cell-intrinsic mobility. This is also in line with the observation that the cells with the highest proliferation rates (LN229, GBM10) in our study are those that show no super-diffusive movement for high cell densities.

The second key aspect we analyzed here is the effect of large cell density gradients on GBM migration, a situation cells potentially face when they reach tumor-stroma boundary, long distance fiber tracts or perivascular spaces. Notably, in the case of GBM cells this boundary is not clear, but a shrinking GBM-cell density with distance from the main tumor mass as well as varying cell densities inside GBM have been reported nonetheless (47). In our experiments subjecting cells to a large-scale cell density gradient, there were two main factors expected to drive the expansion of the clusters: the active forces generated by the cells, transmitted via cell-cell junctions to neighboring cells and a passive expansion driven by increasing pressure due to proliferation far from the cell edge (48). If proliferation and thus a stress gradient is the main driver of expansion, spreading would be expected to have characteristics of a non-active material wetting a substrate and thus velocity and density profiles depend on the distance to the front (48). Given the results for LN229 and GBM10 cells, both dependencies are given, and if those gradients vanish, here achieved via the blockade of proliferation, expansion stops. Consequently, both cell lines appear to expand in a passive way, mostly independent on cell-autonomous processes. In contrast, in U138 cells no consistent cell-density gradient was observed during cluster expansion, independent of proliferation, implying an expansion process driven by active cell migration. Another in silico study also predicted that cellular expansion is mostly growth driven, if the cells have a high proliferation rate but a low mobility (49). Given the determined cell division numbers and single cell migration properties, LN229 and GBM10 cells were both more proliferative and less migratory than U138 and GBM4, further supporting the idea of a mostly passive expansion of LN229 and GBM10.

In the last part of this study, the effect of an increasing cell density was analyzed. In line with the results from the single cell migration experiments and cell exclusion assays, it was found that LN229 and GBM10 but not U138 and GBM4 significantly decrease in their effective mobility, when subjected to high cell densities. One reason for this phenomenon might be their different migratory capacity on the level of individual cells, where LN229 and GBM10 showed markedly reduced mobility and thus high cell densities were more likely to impede their movement. However, even under this assumption, a slowdown and a less super-diffusive movement of U138 and GBM4 cells would be expected too, but especially the super-diffusive movement type remained unaffected. The slowdown and potential migratory arrest of LN229 and GBM10 for high cell densities is in agreement with the expectations from the literature (4–11), arguing that cell density can regulate traction forces (8, 11). Cell density independence of collective migration was reported more rarely, e.g. in a study of mammary carcinoma cells (39) and a previous study of ours in GBM (10). For the motile cells an alignment of cells into parallel strands, combined with a movement in alignment-direction in the forms of small anti-parallel streams was observed. Similarly, streaming behavior was found to be favored before, under conditions of low contractility, high cell mobility and high cell density in epithelial cells and *in silico* (13, 29). These findings provide a potential explanation for the phenotypical differences observed here, as we previously found LN229 cells to have a higher contractility and similar adhesiveness than U138 cells (10). In addition, the observed structures formed by GBM cells resembled those of oncostreams as previously identified to be present in GBM in vivo, promoting collective GBM migration (31). Yet, an *in silico* study on oncostreams predicted anti-parallel streams, as observed here, to be unstable and transform into a parallel streaming motion on the time frame of days (29). Here, we could not observe such a shift in movement pattern but rather found a strengthening of the antiparallel streaming over time. However, there were some factors proposed that block or delay the switch from anti-parallel to parallel streaming: high cell density and cell speeds (29). Given the used modeling parameters, both was given here for the self-organizing cell lines, thus the stability of the anti-parallel streams was to be expected. Of note, those oncostreams were found to resemble the topology of liquid crystals with a quasi long-range order and similar order parameter values as observed here (32). For higher cell densities, an increase was found in the nematic order of the GBM4 and U138 monolayers, accompanied by the expected decrease in the number of topological defects, due to defect annihilation (50–52). In contrast to other studies, we did not observe a favored direction of movement of topological defects relative to their orientation, so they are neither extensile nor contractile (51, 53, 54), albeit the overall movement range of the defects appears to be similar as in epithelial cells (54). Independent on the properties of the nematic defects it appears as though the organization into the liquid crystal-like structure is promoting escape from cell density induced migratory arrest. This interpretation is strengthened by the observation that abrogating the formation of oncostreams blocks tumor infiltration and migration *in vivo* (31). Furthermore, if liquid crystal-like organization is indeed promoting migration under confinement it is expected that the emergence of locally organized clusters promote the migration of those. This situation corresponds to a motile group of cells inside a group of immobile cells. An *in silico* study predicted for this case an impeded flow that eventually leads to the formation of organized, more mobile clusters inside the immobile cell population (29), corresponding to the observations made in LN229 cells here, supporting the notion of self-organization to promote migration.

In conclusion, this study could demonstrate the existence of two migratory distinct phenotypes of GBM cells, one in which cell density reduces effective cell motion and another where cells stay mobile even in very high cell densities. The sustained mobility was independent on proliferation events, but rather dependent on the formation of a local or global cellular organization into liquid crystal-like structures.

**Supplementary Figure 1:** Directionality of individual GBM cells. Box plots show the median (red line), 25 and 75 percentile (box), non-outlier range (whiskers) and outliers (red dots). Sample sizes: n_U138_ = 44; n_GBM4_ = 47; n_LN229_ = 83; n_GBM10_ = 43;

**Supplementary Figure 2:** Manual estimates of the initial cell density at 0 h for all GBM cell lines. Box plots show the median (red line), 25 and 75 percentile (box), non-outlier range (whiskers) and outliers (red dots). Sample sizes: n_U138_ = 184; n_GBM4_ = 139; n_LN229_ = 79; n_GBM10_ = 60;

**Supplementary Figure 3:** Plots of the effect range of a proliferation event on the velocity field as a function of cell density. For each of the four cell lines a plot of the affected range by pre-mitotic contraction in µm (top left) and cells length (top right), as well as the affected range by post-mitotic expansion in µm (bottom left) and cell length (bottom right) is given as a function of effective cell radius. Box plots show the median (red line), 25 and 75 percentile (box), non-outlier range (whiskers) and outliers (red dots). Each boxplot contains at least 70 division events.

**Supplementary Figure 4:** Boxplot of the relative area of the monolayer in which the velocity field is affected by proliferation events. Box plots show the median (red line), 25 and 75 percentile (box), non-outlier range (whiskers) and outliers (red dots). Sample sizes: n_U138_ = 184; n_GBM4_ = 139; n_LN229_ = 79; n_GBM10_ = 60;

**Supplementary Figure 5:** Sample heatmaps of front speeds for the cell exclusion assay. The figure shows examples of the layer speed as a function of time and distance to the cell front for all four GBM cell lines, under control conditions (left column) and when treated with mitomycin C (right column).

**Supplementary Figure 6:** Relation of cell size and expansion rate in cell exclusion assay. A) Graph of the time evolution of average cell size in the field of view as a function of time under control conditions and when treated with mitomycin C. B) to D) Scatter plots of the expansion rate of the layer over the cell density gradient in the layer for U138 cells (B), LN229 (C) and GBM10 (D) cells. Error bars and shaded areas depict the standard error of the mean. Sample Sizes: A) n_U138 CTL_ = 24; n_U138 Mito_ = 11; n_LN229 CTL_ = 29; n_LN229 Mito_ = 15; n_GBM10 CTL_ = 13; n_GBM10 Mito_ = 13 B) n_U138 CTL_ = 24; n_U138 Mito_ = 11 C) n_LN229 CTL_ = 29; n_LN229 Mito_ = 15 D) n_GBM10 CTL_ = 13; n_GBM10 Mito_ = 13.

**Supplementary Figure 7:** Sample images showing different cellular organizations for the four GBM cell lines, at the start of the experiments (left) and after 60 h (right). Please denote the strong emerging order in U138 and GBM4 cells. Scale bar corresponds to 100 µm.

**Supplementary Figure 8:** Analysis of topological defects in U138 cells. A), B) Movement paths of +1/2 (A) and-1/2 (B) defects relative to their orientation. The red line shows the mean movement path over all defects. C), D) Mean squared displacement (C) and the associated scaling coefficient (D) for defects. E) Temporal evolution of the number of defects. Error bars and shaded areas depict the standard error of the mean. Sample size: A)-D) n_+1/2_ = 535; n_-1/2_ = 703; E) n = 184.

**Supplementary Figure 9:** Analysis of topological defects in GBM4 cells. A), B) Movement paths of +1/2 (A) and-1/2 (B) defects relative to their orientation. The red line shows the mean movement path over all defects. C), D) Mean squared displacement (C) and the associated scaling coefficient (D) for defects. E) Temporal evolution of the number of defects. Error bars and shaded areas depict the standard error of the mean. Sample size: A)-D) n_+1/2_ = 279; n_-1/2_ = 286; E) n = 139.

**Supplementary Figure 10:** Analysis of topological defects in LN229 cells. A), B) Movement paths of +1/2 (A) and-1/2 (B) defects relative to their orientation. The red line shows the mean movement path over all defects. C), D) Mean squared displacement (C) and the associated scaling coefficient (D) for defects. E) Temporal evolution of the number of defects. Error bars and shaded areas depict the standard error of the mean. Sample size: A)-D) n_+1/2_ = 603; n_-1/2_ = 685; E) n = 79.

**Supplementary Figure 11:** Analysis of topological defects in GBM10 cells. A), B) Movement paths of +1/2 (A) and-1/2 (B) defects relative to their orientation. The red line shows the mean movement path over all defects. C), D) Mean squared displacement (C) and the associated scaling coefficient (D) for defects. E) Temporal evolution of the number of defects. Error bars and shaded areas depict the standard error of the mean. Sample size: A)-D) n_+1/2_ = 298; n_-1/2_ = 419; E) n = 60.

**Supplementary Figure 12:** Heatmaps of the temporal and spatial evolution of the polar order parameter calculated for the velocity field. Sample sizes: n_U138_ = 184; n_GBM4_ = 139; n_LN229_ = 79; n_GBM10_ = 60.

**Supplementary Figure 13:** Comparison of the speed of ordered and unordered regions in one field of view. A) to D) Ratios of the speed for regions with high nematic order over those with low nematic order for U138 (A), GBM4 (B), LN229 (C) and GBM10 (D). The “-“ sign corresponds to groups with a significant reduction in speed (p<0.05) as determined by the sign test. Box plots show the median (red line), 25 and 75 percentile (box), non-outlier range (whiskers) and outliers (red dots). Sample sizes for t = 0,10,20,30,40,50,60h: B): n_U138_ = 180, 167, 161, 145, 135, 101; n_GBM4_ = 139, 131, 122, 115, 106, 88; n_LN229_ = 79, 79, 78, 78, 77, 73;n_GBM10_ = 32, 27, 28, 29, 28, 21.

**Supplementary Video 1:** Cell exclusion assay for GBM4 cells, showing expansion and signs of self-organization.

**Supplementary Video 2:** Cell exclusion assay for U138 cells, showing expansion and signs of self-organization.

**Supplementary Video 3:** Cell exclusion assay for GBM10 cells, showing expansion and no signs of self-organization.

**Supplementary Video 4:** Cell exclusion assay for LN229 cells, showing expansion and no signs of self-organization.

**Supplementary Video 5:** Cell exclusion assay for LN229 cells treated with mitomycin C, showing expansion and no signs of self-organization.

**Supplementary Video 6:** Confluent monolayer for GBM4 cells, showing migration and emergence of anti-parallel moving streams and self-organization.

**Supplementary Video 7:** Confluent monolayer for U138 cells, showing migration and emergence of anti-parallel moving streams and self-organization.

**Supplementary Video 8:** Confluent monolayer for GBM10 cells, showing little migration and no emergence of anti-parallel moving streams or self-organization.

**Supplementary Video 9:** Confluent monolayer for LN229 cells, showing migration and no large-scale emergence of anti-parallel moving streams or self-organization.

**Supplementary Table 1:** Number of detected cell division events and number of analyzed fields of view for each cell line.

